# Creating anatomically-derived, standardized, customizable, and three-dimensional printable head caps for functional neuroimaging

**DOI:** 10.1101/2024.08.30.610386

**Authors:** Ashlyn McCann, Edward Xu, Fan-Yu Yen, Noah Joseph, Qianqian Fang

## Abstract

**Significance:** Consistent and accurate probe placement is a crucial step towards enhancing the reproducibility of longitudinal and group-based functional neuroimaging studies. While the selection of headgear is central to these efforts, there does not currently exist a standardized design that can accommodate diverse probe configurations and experimental procedures.

**Aim:** We aim to provide the community with an open-source software pipeline for conveniently creating low-cost, 3-D printable neuroimaging head caps with anatomically significant landmarks integrated into the structure of the cap.

**Approach:** We utilize our advanced 3-D head mesh generation toolbox and 10-20 head landmark calculations to quickly convert a subject’s anatomical scan or an atlas into a 3-D printable head cap model. The 3-D modeling environment of the open-source Blender platform permits advanced mesh processing features to customize the cap. The design process is streamlined into a Blender add-on named “NeuroCaptain”.

**Results:** Using the intuitive user interface, we create various head cap models using brain atlases, and share those with the community. The resulting mesh-based head cap designs are readily 3-D printable using off-the-shelf printers and filaments while accurately preserving the head topology and landmarks.

**Conclusions:** The methods developed in this work result in a widely accessible tool for community members to design, customize and fabricate caps that incorporate anatomically derived landmarks. This not only permits personalized head cap designs to achieve improved accuracy, but also offers an open platform for the community to propose standardizable head caps to facilitate multi-centered data collection and sharing.

## 1 Introduction

Functional brain imaging techniques play essential roles in our quest for a better understanding of complex human brain activities. Traditional neuroimaging studies utilizing functional magnetic resonance imaging (fMRI) paradigms have been largely confined to laboratory or clinical settings, mostly due to the immobility of fMRI instruments.^1^ In recent years, the push towards understanding human brains in natural environments,^2, 3^ including during day-to-day tasks and social interactions,^4^ demands lightweight, wearable,^5–7^ and flexible neuroimaging modalities as well as robust data acquisition for long-term monitoring and recording.^8^

Functional near-infrared spectroscopy (fNIRS) has rapidly emerged as a competitive option for fulfilling the needs of mobile and long-term neuromonitoring,^8^ offering compact and wearable form factors, relatively low cost, and safety for long-time measurements using non-ionizing near-infrared light.^9^ Compared to electroencephalography (EEG), another widely used neuroimaging modality that also offers portability and wearability,^10^ fNIRS^6, 11, 12^ provides improved spa-tial resolution and a rich set of physiological measurements, including oxy-hemoglobin, deoxy-hemoglobin, oxygen saturation, blood volume, among others.^13^ Clinical and research applications of fNIRS include brain-injury monitoring,^13^ investigating various psychiatric and neurological dis-orders,^14, 15^ as well as real-time monitoring of neural activation in response to specific tasks and conditions.^16–18^

Similar to EEG, many fNIRS studies use wearable head caps to attach optical probes – an assembly containing optical sources and detectors with driving electronics and light-coupling fibers – securely to the subject’s scalp to enable robust data acquisition.^19^ The head cap plays several important roles in fNIRS/EEG experiments and directly impacts the quality of the acquired data.^20^ First, it is often used as a mounting frame to firmly attach the optodes to the subject’s head, and stabilize the optode-to-scalp coupling by applying gentle pressure through the elasticity of the cap and the built-in straps.^21^ Secondly, it often provides standardized head landmarks in terms of pre-fabricated sockets or markings aligned with EEG 10-20/10-10/10-5 systems^22, 23^ to guide the placement of optodes above the desired brain functional regions.^24^ Thirdly, fNIRS head caps are often made of light-absorbing materials, shielding the light-sensitive optical sensors from ambient and stray light and increasing the robustness of optical measurements.^25^ Although many fNIRS head caps have been derived and modified from EEG head cap designs,^26^ they vary dramatically in terms of materials, sizes, thicknesses, head-landmark consistencies, mounting grommet shapes, density, spatial arrangements etc, between different studies. Building standardized fNIRS head caps that can accommodate diverse probe designs has been discussed recently within the fNIRS community,^27^ especially recognizing the importance of using such standardized head caps in cross-instrument validations and group-based data analyses.

Developing a standardized head cap must address several practical challenges. First, the sizes and shapes of optical probes vary greatly,^26^ depending on applications and cost constraints. For example, some fNIRS studies use small compact fNIRS probes^28–30^ while others use full-head high-density probes.^5, 31^ Secondly, 10-20 landmark locations are often manually measured and marked to account for variations in head size and shape, which results in relatively lengthy setup process as well as inconsistencies in probe placement due to operator variability.^32^ The current alternative is to use a generic EEG cap, while simplifying the donning process by guiding probe placement, it also introduces variability in probe placement.^33^ Multiple studies have concluded that incorporating subjects’ anatomical information, as well as consistent probe placement directly impacts fNIRS reproducibility^34^ and group-level analyses,^35^ stressing the importance of incorporating anatomically relevant landmarks into the cap design. While using the 10-20 system does not completely eliminate ambiguities and operator-dependent variations, including uncertainties associated with selection of cranial reference points,^23, 32^ it is nonetheless one of the most commonly adopted approaches to provide consistent spatial reference points for head-based probe placement,^36^ either through 10-20 or study-specific systems^37^. Thirdly, hair is a major confounding factor for obtaining robust fNIRS measurements.^38^ It is important that the head cap provides ample space for accessing the hair near the optodes and allows easy adjustments. Additionally, fNIRS probe setup often takes significant effort and time, ranging from a few minutes to just under an hour^39–41^ before the experiment to ensure good signal quality. A lightweight, easy-to-mount head cap with built-in 10-20 landmark positioning guidance could greatly accelerate this process.

It is generally agreed upon that an ideal fNIRS head cap should achieve 1) stabilization of optodes, 2) adequate user comfort, and 3) ease of use and implementation.^20^ Compatibility with EEG is another factor that has been considered, as many previously reported caps have been designed to permit simultaneous measurement of EEG and fNIRS data.^11, 41^ In addition, while there is a variety of head caps available commercially, many of these products are specifically tailored for use with particular commercial fNIRS/EEG systems and may not be readily applicable to other fNIRS systems without significant modification. Typically, they offer limited options that can account for differences in head sizes, head shapes, and optode locations.^20, 42^ Recent studies have investigated three-dimensional (3-D) printed caps, which provide benefits in flexible customization, easy cross-lab reproducibility, and easy fabrication.^43^ However, there is currently no publicly available software or head cap models to design and fabricate 3-D printable neuroimaging caps. More specifically, there is a lack of software tools for designing anatomically derived head caps.

Here, we introduce an open-source software and workflow for designing and creating anatomically derived, personalized, and 3-D printable neuroimaging head caps that could potentially address the needs of standardizing fNIRS measurements, cross-lab reproducibility, flexible customization for diverse optode designs, and ease of fabrication, as well as various ergonomic considerations. This workflow has been implemented as an add-on to Blender, a powerful and widely used open-source 3-D modeling platform.^44^ Through an intuitive graphical user interface (GUI), this fNIRS/EEG head cap design add-on, named “NeuroCaptain”, allows users to conveniently extract head surface mesh models from volumetric magnetic resonance imaging (MRI) scans or brain atlases, compute anatomically derived 10-20 landmark positions, customize landmark grommet shapes and sizes, and convert wireframe head cap mesh models into water-tight 3-D printable models. NeuroCaptain seamlessly integrates our high-quality open-source MATLAB-based brain mesh generation toolbox, Brain2Mesh,^45^ with the rich 3-D modeling functionalities offered by Blender, making it possible to widely disseminate among the community and permit researchers to create reproducible head caps across institutions and studies. We want to highlight the hypothesis we aim to test with this work is centered around the idea that a publicly accessible, anatomically-derived cap design and fabrication pipeline can help the community reinforce optode placement reproducibility across diverse study populations and accommodate wide varieties of instrumentation designs. Characterizing the functionality of the cap in experimental settings, such as robustness of coupling, head shape conformity, and user comfort, is of great interest, these factors will be evaluated in a future study.

In the following Methods section, we will detail the workflow for 3-D head surface mesh generation, 10-20 point calculations, and steps for creating water-tight 3-D printable head cap models. In the Results section, we show examples of printed head caps with various sizes and mesh densities. We also validate the accuracy of the printed head caps by measuring the head landmark locations over a sample cap worn by a healthy volunteer.

## 2 Methods

### 2.1 Overall workflow

The goal of this work is to develop a versatile and easy-to-use software tool that can create 3-D printable head cap models derived from anatomical imaging, such as brain atlases or subject-derived MRI scans, with embedded standardized head landmarks to guide probe placement. The workflow of the data processing is illustrated in Fig. 1. We divide the overall workflow into 3 sections: 1) creating the head surface model, 2) calculating 10-20 landmarks and generating the cap, and 3) making the cap model 3-D printable. In the following sections, we detail each of the steps. The entire workflow is built upon open-source tools and libraries with a potential for broad dissemination.

**Fig 1.**
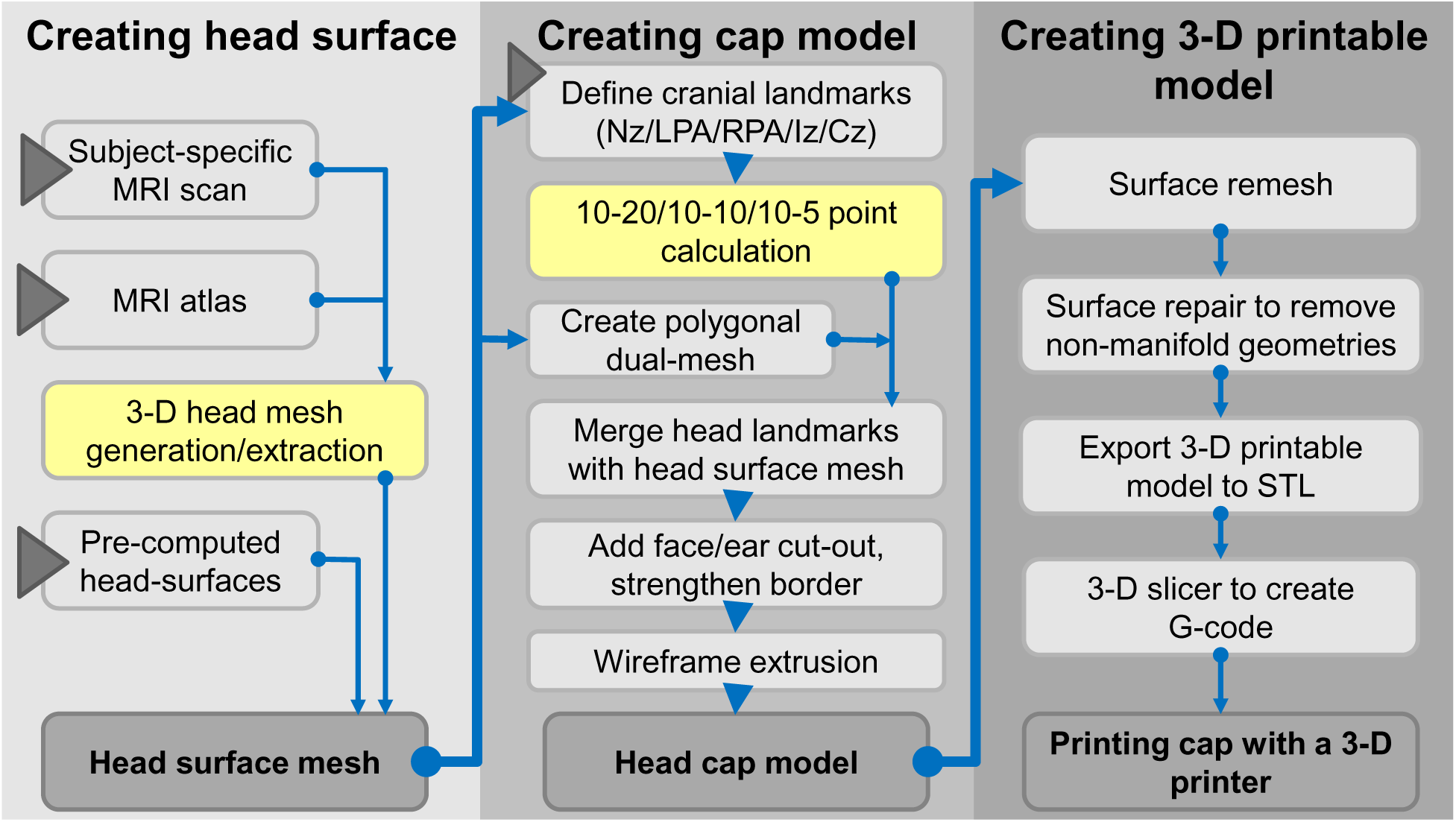
General workflow of the NeuroCaptain cap design pipeline. Solid triangles indicate user inputs; yellow-shaded blocks are computed inside MATLAB/Octave.

### 2.2 Creating head surface

The first step of creating a 3-D printable head cap is to generate a triangular surface mesh model representing a human head. There are several neuroanatomical analysis tools that can create scalp surface mesh, including FMRIB Software Library(FSL),^46^ BrainSuite^47^ and our in-lab software, Brain2Mesh.^45^ If such a surface model has been previously generated, a user can directly import this mesh model into Blender and move on to the next step. Many commonly used surface mesh formats are supported in Blender, such as standard triangle language (STL) format, object file format (.off), or Wavefront format (.obj), among others. Moreover, we also support a newly de-veloped JavaScript Object Notation (JSON) based JMesh format^44^ by our group to facilitate shape data exchange.

In addition, generating subject-specific head caps using a subject’s own imaging data may improve overall anatomical accuracy^48, 49^ and subsequently, improved cap-scalp conformity. However, subject-specific imaging data may not always be available. In such a case, NeuroCaptain allows users to import a segmented atlas model. In this case, we apply Brain2Mesh to automatically extract the scalp/head surface by setting a threshold in an MRI scan, multi-label, or probabilistic segmentation of the head. As we described in,^45^ Brain2Mesh employs the *ɛ*-sampling algorithm^50^ to create high-quality surface meshes from gray-scale or multi-labeled 3-D volumetric images. The mesh density is controlled by setting the maximum radius of the circumscribing circle of the triangles on the surface. The computation of the *ɛ*-sampling algorithm is relatively fast and only takes a few seconds to complete.

### 2.3 Creating cap model

A notable feature of our 3-D printable cap design is the built-in 10-20 landmarks with anatomically determined locations. To achieve this, we have developed a semi-automated algorithm to compute the 10-20 head landmark 3-D positions over a triangulated head surface. This algorithm requires a user to first manually specify 5 key cranial landmarks, including nasion (Nz), left preauricular point (LPA), right preauricular point (RPA), inion (Iz), and an initial estimation of the vertex (Cz0). Using LPA, RPA, and Cz0 positions, we compute the coronal reference cross-sectional curve (a loop) of the head surface mesh (assumed to be a closed surface with a single compartment). We then traverse from LPA towards Cz0 and further to RPA to identify the portion of the closed loop over the top of the head surface and calculate the curve distance between LPA-Cz0-RPA. Similarly, using Nz, Iz, and Cz0, we compute the sagittal reference cross-section and return the curve distance between Nz-to-Cz0 and Iz-to-Cz0. Because the true vertex position (Cz) is supposed to bisect both sagittal and coronal reference curves, we use an iterative algorithm to update Cz, starting from Cz0, until the distance between subsequent estimates drops below a predefined threshold (10*^−^*^6^ in mesh coordinate unit). As a result, this algorithm outputs a list of 10-20 (or 10-10 or 10-5) landmark 3-D positions defined over the head surface mesh space (see Fig. 2 top-middle panel). A Delaunay triangulation is implemented to generate a triangular mesh, with each node corresponding to a 10-20 coordinate (see Fig. 2 top-right panel), to facilitate the import of these positions into Blender.

**Fig 2.**
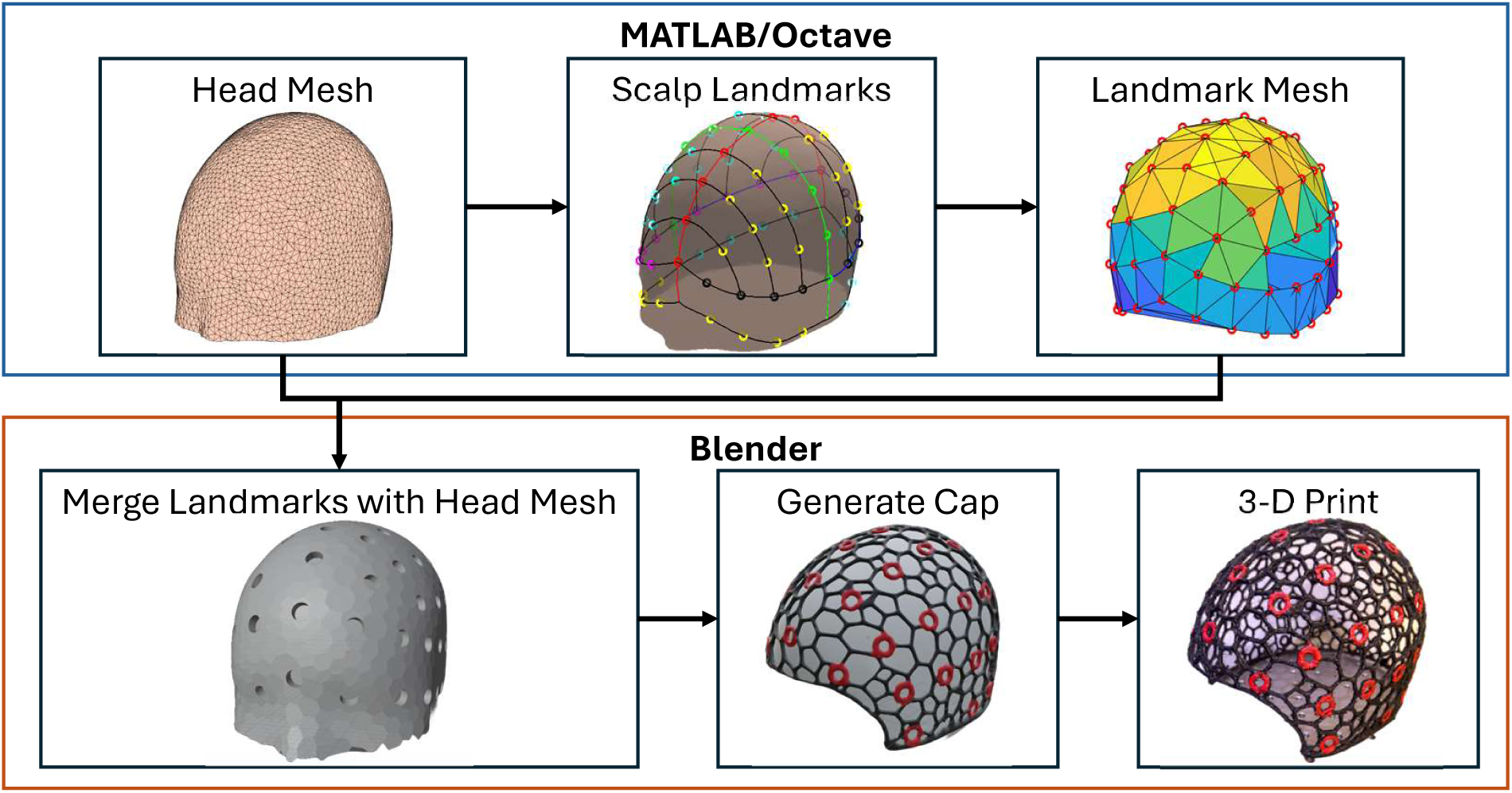
Diagram showing the process of incorporating 10-20 landmarks in a 3-D printable cap model. The top half shows the steps computed in MATLAB/Octave; the bottom half shows the outputs in Blender.

It is important to note that the difficulty in accurately selecting the cranial landmarks is an inherent challenge when using the 10-20 system,^23^ particularly the selection of the inion location.^51^ However, such a challenge is not unique to the cap design method proposed in this work. As an alternative to manually selecting cranial landmarks on the head surface, researchers can import 3-D coordinates of cranial reference points or optode positions directly obtained by measurements, such as those acquired through 3-D digitization or photogrammetry.^49^ Through the support of user-defined grommet locations, a user can gain significant flexibility in customizing the 3-D printable cap to accommodate diverse experimental needs in measurement locations, areas, or mounting of the optical probes.

We also pre-process the head surface mesh before merging with the 10-20 position markers. We apply a dual-mesh transform using a built-in Blender add-on named “tissue tools” to convert the triangular head mesh to a polygonal mesh. The resulting mesh is courser while still adequately preserving surface topology. Once the mesh is converted to a wire-frame model, this feature will allow easy access to underlying hair and fiber couplers for on-the-spot adjustments during experi-ments.

In the next step, we integrate the anatomical landmarks into the previously extracted head surface. This is achieved in three main steps: 1) registering the 10-20 landmarks to the head surface, 2) creating an array of “landmark grommets” in the form of user-specified shapes centered at each 10-20 location oriented along the surface normal direction, and 3) performing a Boolean operation to modify the head surface to embed the landmark grommets. This results in a surface mesh with holes in the shapes of user-selected marker outlines at the location of all head landmarks (see Fig. 2 bottom-left panel). To automate this process, we have developed a “geometry-node” based procedural modeling program in Blender, allowing users to perform this multi-step operation using a single button-click. The geometry node program has various parameters, including the size of the landmark grommets, that can be adjusted and fine-tuned to suite diverse needs. This landmark embedding operation can be applied multiple times, allowing users to incorporate not only head landmarks, but also specific fiber/optode mounting sockets of different sizes to the head cap to accommodate different probes.

The detailed steps of the geometry nodes procedural modeling program is further visualized in Fig. 3. First, for every 10-20 position, the closest location in the input head surface is calculated (using the “Geometry Proximity” geometry node module) and used for translating the input 10-20 mesh to spatially align with the head mesh (“Set Position”). As a separate step, the landmark mesh, as shown in Fig. 2 top-right, is oriented with the head surface at each 10-20 point vertex. To allow consistent alignment, the normals on the surface triangles of the head mesh are calculated (“Sample ‘normals’ Nearest Surface”). The resulting set of vectors is used to rotate the landmark grommet shapes to be orthogonal to the head surface normal direction (“Align Euler to Vector”). The geometry node program iterates over each 10-20 position and repeat the above transformation, resulting in an array of normal-aligned landmark grommets placed over each vertex in the 10-20 mesh (“Instance on Points”). Once the landmark grommet array is created, a mesh “Boolean difference” operation is executed, subtracting the landmark grommet array from the head mesh. The resulting output is a head mesh with distinct holes at each 10-20 location, as visualized in Fig. 2 bottom-left.

**Fig 3.**
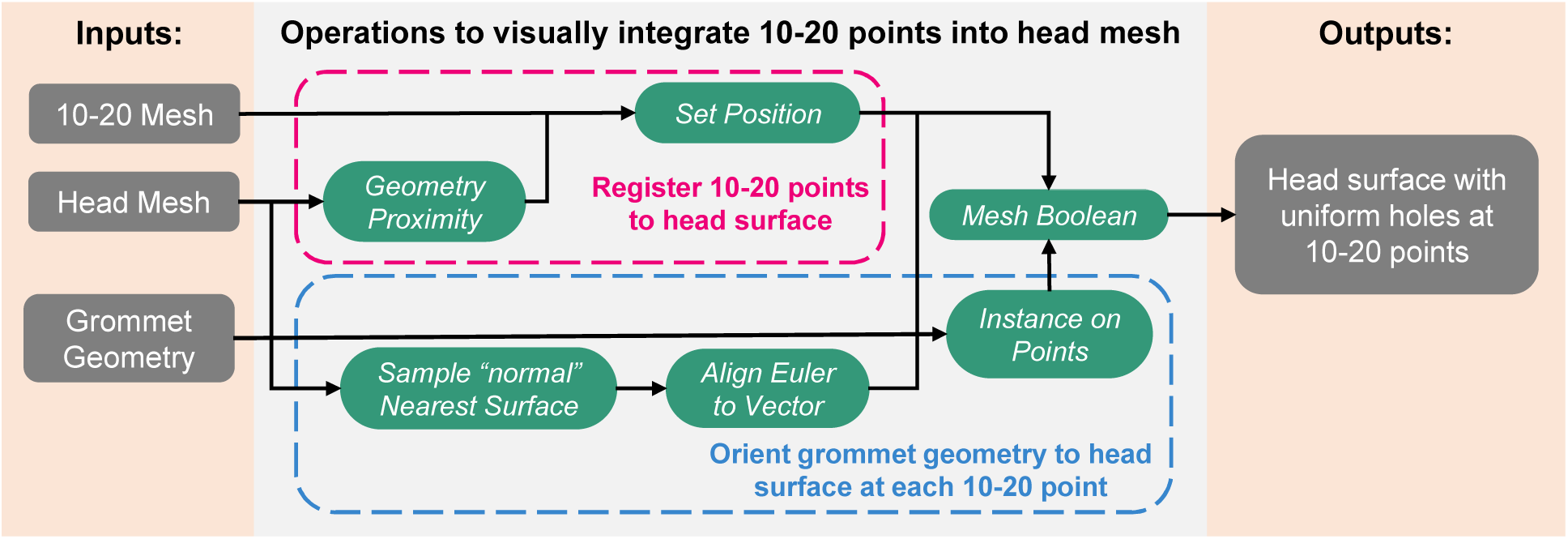
Workflow diagram illustrating the “geometry node” programming steps in Blender to convert 10-20 positions to surface grommets. The green-colored blocks indicate Blender geometry node functions. Overarching steps are specified by dotted lines encompassing the nodes.

Additional steps are performed to create the proper cap margin shape for ease of wearing. A “Boolean difference” is made first between the head mesh with a second box-shaped mesh (“face-box”) to create the opening over the face. Another “Boolean difference” is subsequently performed between the remaining head mesh and a box-shaped mesh (“neck-box”) to set the lower boundary of the cap towards the neck. Nz guides the placement of the face-box. The vertex corresponding to Nz is selected such that the center point of the face-box is aligned with the coordinate of Nz. The width of the face-box is set as 1/3 of the width of the head model. The neck-box is set to have the full width of the head model along the *x* and *y* dimensions, with the *z* dimension length setting to 1/3 of head model. Optional Boolean cuts can be made to shape the margin of the cap to create comfortable placement over ears. Once achieving the desired margin shapes, the edges of the mesh are transformed into a 4-sided polygon (known as a “quad”), such that the cap becomes an open wireframe mesh. Finally, the edges along the exterior margin of the cap are thickened to protect the cap from tearing.

To complete the head cap model, we extrude all edges of the polyhedral head cap mesh by a user-defined wire thickness using a transform known as “wireframe” in Blender. This converts the cap mesh into a closed wireframe surface mesh, getting ready for preparing the 3-D printable model in the next step.

### 2.4 Interfacing with optical sources and detectors

We want to highlight that the “grommet” structures incorporated into the cap design in Fig. 3 are not restricted to 10-20 landmarks. When desired, they can be placed underneath the target source/detector locations, with shapes/sizes that can facilitate the convenient mounting and securing of the optodes. It has been a widely utilized practice in 3-D printing to design and print models with interfaces that allow easy interconnection with non-3-D printed parts, via common mechanisms such as pressure-fitted, notched, geared or threaded structures built-in to the 3-D model of the printed components. With a printing nozzle size of 0.1 – 0.2 mm, 3-D printed parts are well capable of creating mechanical interfaces with size-sensitive parts such as optical couplers or modules. This will require the cap designer to utilize Blender’s modeling capabilities that are outside of those streamlined by NeuroCaptain to design and incorporate these mechanisms into the cap model before fabrication.

For simple fNIRS fiber/coupler based interfaces, NeuroCaptain printed caps can use the default circular grommets for connecting with optodes. When printed with the right inner-diameter size, these grommets can be used to directly insert/secure optical fiber based sensors over the cap. If the accuracy of the 3-D printed grommet is not sufficient, the wireframe based grommet can also be used to embed commercially available rubber grommets commonly used in EEG/fNIRS head cap designs.^43^

### 2.5 Creating 3-D printable model

Once the cap model is completed, we then start the creation of the final 3-D printable model (see Fig. 1 right column). A 3-D printable model must be a water-tight closed surface without topological defects. The repeated mesh manipulations applied in previous steps, such as Boolean operations and wireframe extrusion, introduce non-manifold and self-intersecting triangles. To mitigate these defects, we apply a “remesh” step to re-tessellate the extruded wireframe space to fix the surface defects, followed by a mesh simplification step via merging vertices within a pre-set distance threshold. Additional mesh repairing and cleaning operations are manually performed, including recalculating normals, filling holes, and deleting zero-area faces and zero-length edges. Finally, the head cap model is exported from Blender as a standard triangle language (STL) file, which is compatible with most 3-D printing software for generating 3-D printer hardware instructions, known as the G-code.

### 2.6 Graphical user interface design and automation

The complex and multi-step neuroimaging cap generation workflow, as depicted in Fig. 1, has been fully automated using Python programming and integrated with the Blender graphical environments in the form of an add-on, named “NeuroCaptain”. Using Blender’s Python programming interface (i.e. the `bpy` module), we have created an intuitive graphical user interface (GUI) to allow users perform each of the steps using simple button clicks.

In Fig. 4, we show a diagram describing the data communication between Blender and Octave/MATLAB programming environments using a combination of Python and MATLAB scripts and toolboxes. Specifically, the vertices, triangular faces and cranial landmark information are extracted from Blender and written into a JSON-based data exchange file using Python; by calling Octave inside Python, we can compute the 10-20 landmark positions using the `brain1020` func-tion in the Brain2Mesh toolbox. The computed 10-20 positions are tessellated and returned back to Blender using another JSON based data file.

**Fig 4.**
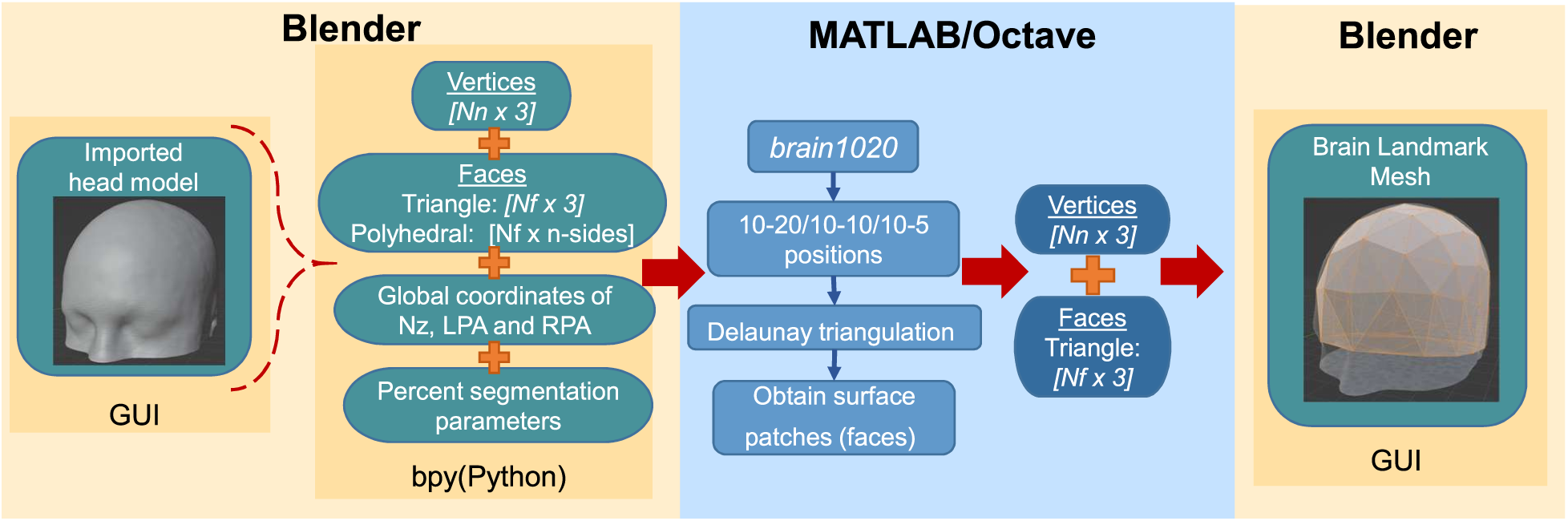
Diagram visualizing the data exchange between Blender and MATLAB/Octave environments, using a combination of Blender-Python application programming interface (API) and external MATLAB toolboxes (including Iso2Mesh, Brain2Mesh etc).

The GUI design reflects the methodology of generating the head caps, with each step automated and easily accessible through simple user interactions. Fig. 5 shows a screen capture of the NeuroCaptain add-on interface in the Blender environment. The functions exposed in the NeuroCaptain add-on panel are roughly organized in the same order as the overall workflow shown in Fig. 2. For each step, user customization is permitted by popping up an option dialog as shown in the middle of Fig. 5. Using these option dialog, users can easily set head landmark density (10-20, 10-10 and 10-5), the density of cap meshing, the size and shape of the landmark socket, and the thickness of the wireframe for the cap. Users can also apply linear scaling to any generated cap mesh model using Blender’s built-in “Scale transformation”. This may be necessary for scaling an atlas to a subject’s head size or creating a range of sizes from one atlas. We also provide a circumference estimator allowing users to measure the circumference of the head surface used for creating the cap, so that a designer can utilize this measurement to scale the cap to fit any target head size. To demonstrate the ease-of-use of our cap design software, we provide a brief video tutorial, named “Visualization 1” shown as a link in the caption of Fig. 5.

**Fig 5.**
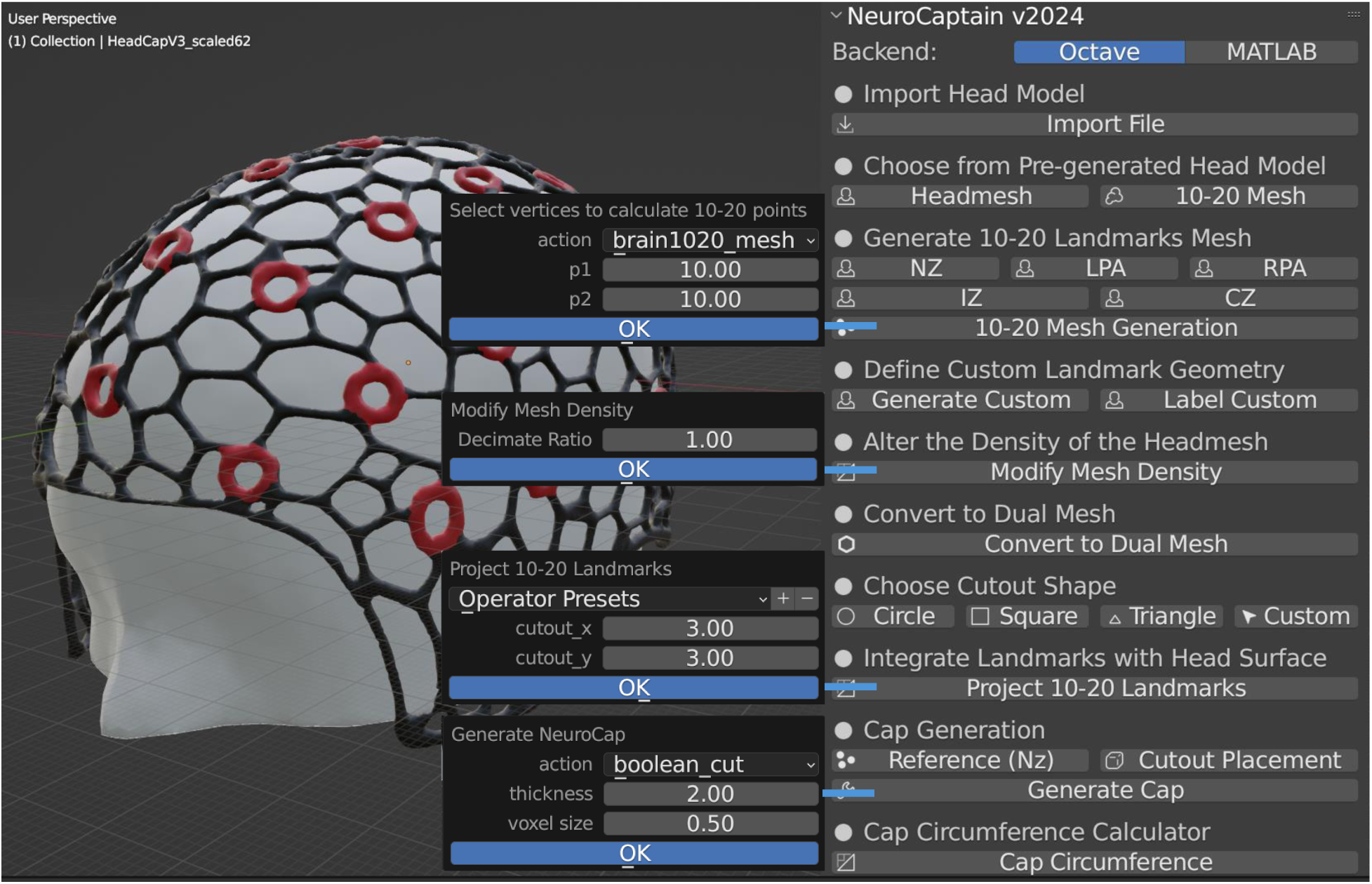
Screenshot showing NeuroCaptain graphical user interface (GUI) in Blender. Parameter dialog windows are shown in the middle allowing users to customize diverse cap setting. An animation showing the cap design process using NeuroCaptain can be accessed as Visualization 1.

## 3 Results

### 3.1 Head cap customization and standardized head cap library

The highly flexible and versatile head cap design workflow described above can be tailored to specific experimental needs and applications while the built-in atlases and anatomically derived landmarks also offer a standardizable framework for creating head caps that can be reproduced and compared across research labs and studies. Our open-source software, NeuroCaptain, allows users with minimal experience to customize an array of cap parameters. All head caps presented in this work are generated using the open-source GNU Octave platform as the back-end, further highlighting the wide accessibility of this tool. To further facilitate adoption, we have also created a library of pre-generated 3-D printable head cap models. This library includes head caps derived from the widely used Colin27 atlas and the Neurodevelopmental MRI atlas (NDMRI) library, with age-dependent atlases ranging between 6 months and 84 years old. To demonstrate the varieties of features and customization options that NeuroCaptain supports, in Fig. 6, we show a few sample cap designs from this cap library.

**Fig 6.**
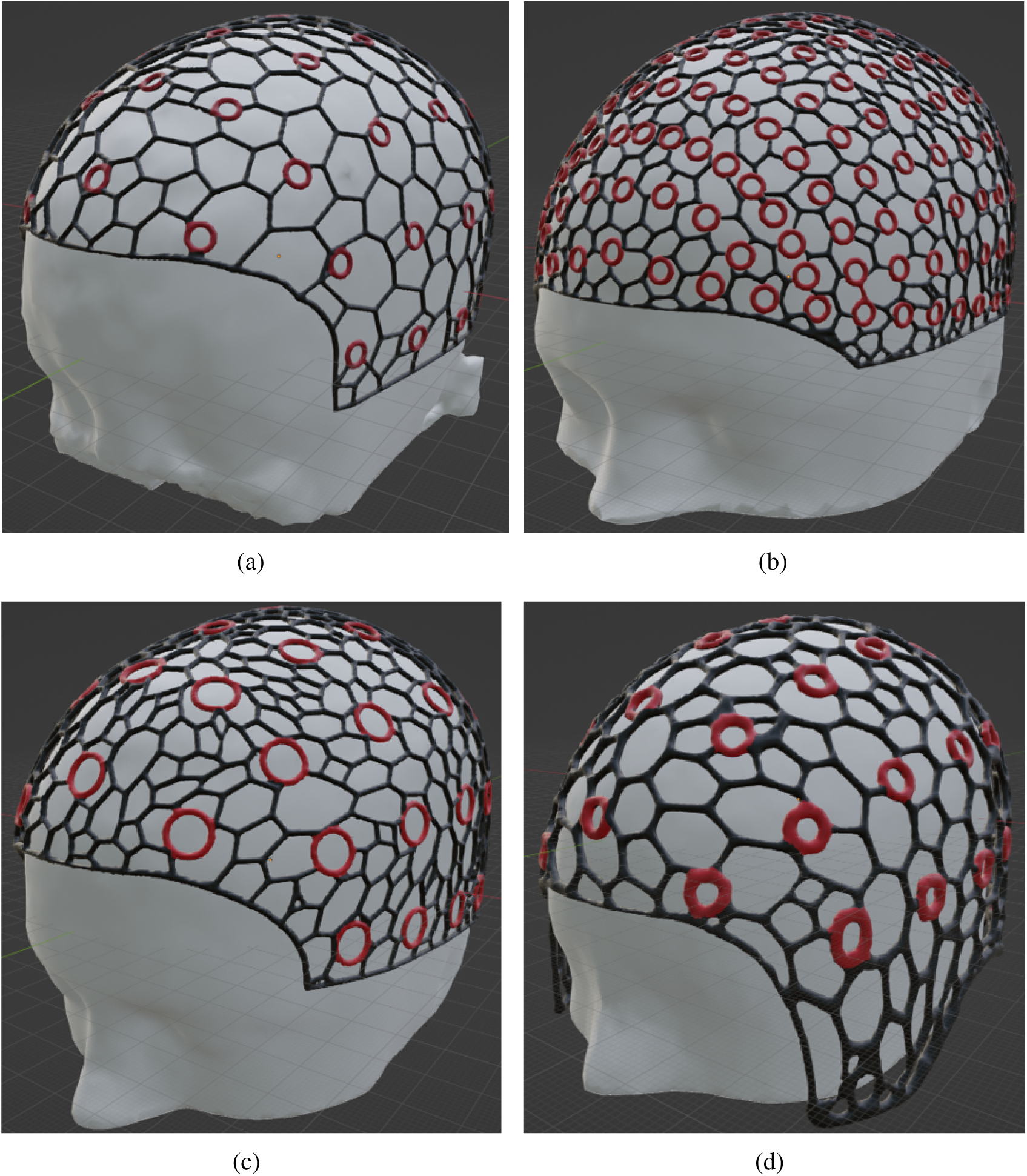
Sample head cap models generated by NeuroCaptain, including caps derived from a) a 2-year-old atlas with coarse wireframe, b) a 35-39 year-old atlas with dense wireframe and 10-5 grommets, c) an 80-84 year-old atlas with large diameter 10-10 markers, and d) a 40-44 year-old atlas with thick wireframe and ear cut-out designs. The red circles denote the 10-20 landmarks embedded in the cap design.

Fig. 6(a) shows a coarse wireframe head cap design with a decimated mesh using a “keep-ratio” of 0.05 (i.e. keeping 5% of the edges from the original imported head surface) and a wireframe with a 2 mm thickness, derived based on the 2-year-old atlas from the NDMRI library. The locations of the 10-10 head landmarks are computed and embedded in this cap with a circular grommet of 4 mm radius. The resulting cap has a circumference of 50.2 cm. In Fig. 6(b), we show a dense cap design with a keep-ratio of 0.1, a circumference of 56.0 cm, and wireframe thickness of 2.5 mm, embedded with 10-5 landmarks derived from the 35-39 year-old atlas in the NDMRI library using a circular grommet with a 3.5 mm radius. In Fig. 6(c), we show another cap derived from an 80-84 year-old NDMRI atlas with large (6 mm radius) 10-10 grommet and a circumference of 57.4 cm. In all of the 3 aforementioned head caps, a flushed bottom edge design is used. Finally, in Fig. 6(d), we show that one can also customize the shape of the bottom edge of the cap with an elongated preauricular section and a loop-cut at the location of the ears. Thicker (3.5 mm diameter) wireframe designs are also used for both the head mesh as well as the circular 10-10 landmark grommets. This cap has a circumference of 55.3 cm. Our caps are printed with wireframe thicknesses ranging from 2.5 mm to 3.5 mm. The desired thickness often depends on the user’s needs and the stiffness rating of the printing filament. Generally, a thinner wireframe provides more elasticity and flexibility but are more prone to breaking. A thicker wireframe offers stronger structural integrity at a cost of reduced stretchability. Our recommended range of 2.5 mm to 3.5 mm wireframe size demonstrates good mechanical robustness even after repeated use.

### 3.2 Fabrication of head caps with 3-D printing

To validate our head cap design workflow, we have fabricated a number of fNIRS head cap using various design parameters and 3-D printing techniques. In the remaining section, we describe detailed processes of printing our designed head caps using multi-filament and single-filament 3-D printers. The 3-D wireframe caps shown in Figs. 7(a-b) are fabricated by a commercial Stratasys F170 3-D printer (Stratasys, Eden Prairie, USA) using a thermoplastic polyurethane (TPU) filament (FDM TPU-92A, Stratasys, Eden Prairie, USA) to print the cap body and a quick support release (QSR), water soluble filament (FDM QSR Support 60ci, Stratasys, Eden Prairie, USA) as the supporting materials. The NeuroCaptain-generated STL file is imported into a proprietary Stratsys slicer software: GrabCAD Print. In this software, the cap is first reoriented such that the open-end of the cap is facing upwards, and the apex of the cap is on the plate of the machine to minimize support material use and printing time. A rendering of the sliced model in GrabCAD Print, showing supporting materials in orange and TPU in green, can be found in Fig. 7(a). After generating the G-code and sending the sliced model to the printer, this cap takes about 40-48 hours to print. The final print consumes 105.06 grams of TPU materials and 323.01 grams of the supporting materials. A photo of the partially printed cap on the Stratasys printer is shown in Fig. 7(b). The completed print is then submerged in a support cleaning apparatus (SCA 1200HT) station to dissolve the support materials for at least 8 hours at a solution temperature of 70*^◦^* C. The solu-tion to dissolve supporting materials consists of water and 6 packets of Ecoworks cleaning agent (Stratasys, Eden Prairie, USA). The caps then go through minor post-processing steps to clean up the print including a rinse with water and removing excess TPU threads. The cost of the raw materials needed for this print is about $58 USD. The overall cost amounts to $155 USD when including additional labor costs of outsourcing fabrication.

**Fig 7.**
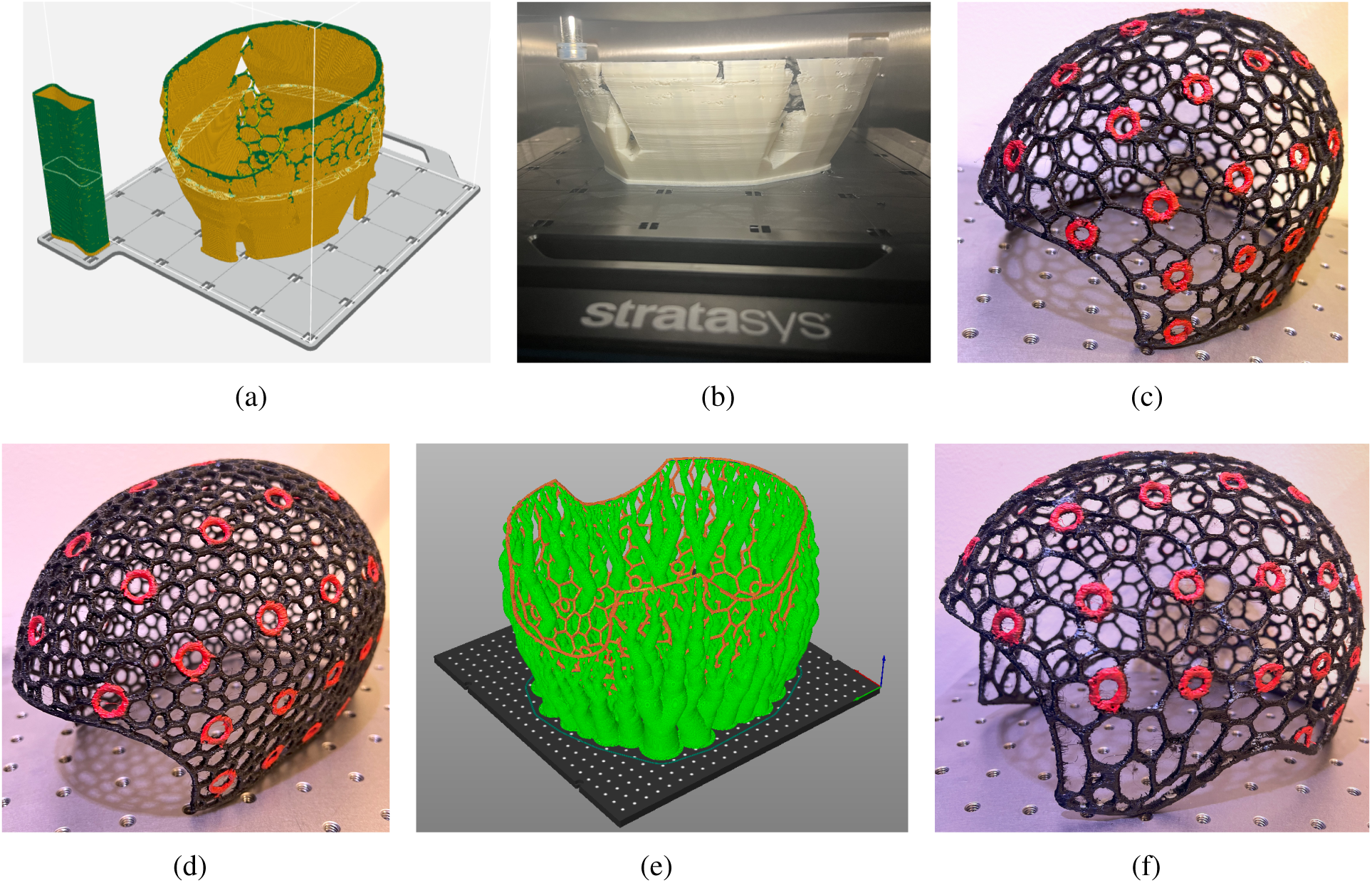
Sample 3-D printed caps. We first show the sliced cap model in (a) prepared for a multi-filament printer (Stratasys F170), with supporting materials shown in orange and thermoplastic polyurethane (TPU) cap shown in green. A photo of the partially printed cap on the Stratasys printer is shown in (b), and the completed cap after post-processing is shown in (c). A similar but dense wireframe cap was also printed and shown in (d). In (e), we show the sliced model prepared for a single-filament printer (Voron 2.4) with organic supports. Green indicates the organic support and orange indicates the actual cap, both printed with the same TPU material. The completed cap is shown in (f) with ear cut-outs. The 10-10 landmarks on all printed caps are painted red with acrylic paint to provide better visual guidance.

In Fig. 7, we give an example of fabricated head caps using a low-cost, large-format 3-D printer based on a single-filament Voron 2.4 (VORON Design) printer with only TPU-95A fila-ments (Cheetah, NinjaTek, Lititz, USA). The slice thickness of the print is set to 0.2 mm. An open-source slicer software, “PrusaSlicer”, is used to generate the G-code for this printer. To utilize a single filament to print the full cap with hanging structures, a special support structure, known as organic supports, is used, shown in green in Fig. 7(e). The completed cap is shown in Fig. 7. We want to note that the organic support is easy to detach from the main print without causing damage to the print. Compared to the water-soluble supports of the previous print, the organic supports are cheaper and requires almost no additional time for post-processing compared to over 8 hours post-processing time when using the water-soluble support. For the sample cap shown in Fig. 7(e), the total print consumes 358 grams of TPU-95A filaments and roughly 16 hours of printing time. The materials cost for this print is $31.36 USD, making the total fabrication cost to roughly $135 USD with the addition of labor cost.

### 3.3 Head cap landmark validation

#### 3.3.1 Validation using 3-D printed head model

We first validate the accuracy and reproducibility of the 10-20 landmark grommets positions using a 3-D printed head model and a size-matched cap derived from the same head surface. The head model used in this benchmark is the NDMRI atlas for ages 40-44. Both the head surface and the cap are scaled at the same circumference of 64 cm. On the head surface model, the target 10-10 positions are indicated with circular through-holes. The head model is printed on an Ender-5 (Creality, China) printer using white PLA filaments. The entire model took around 16 hours to complete, with the resulting head model print shown in Fig. 8(a). The cap was printed using the Voron 2.4 (VORON Design) printer with a TPU-95A filament, fabricated about 14 months prior to the printing of the head model.

**Fig 8.**
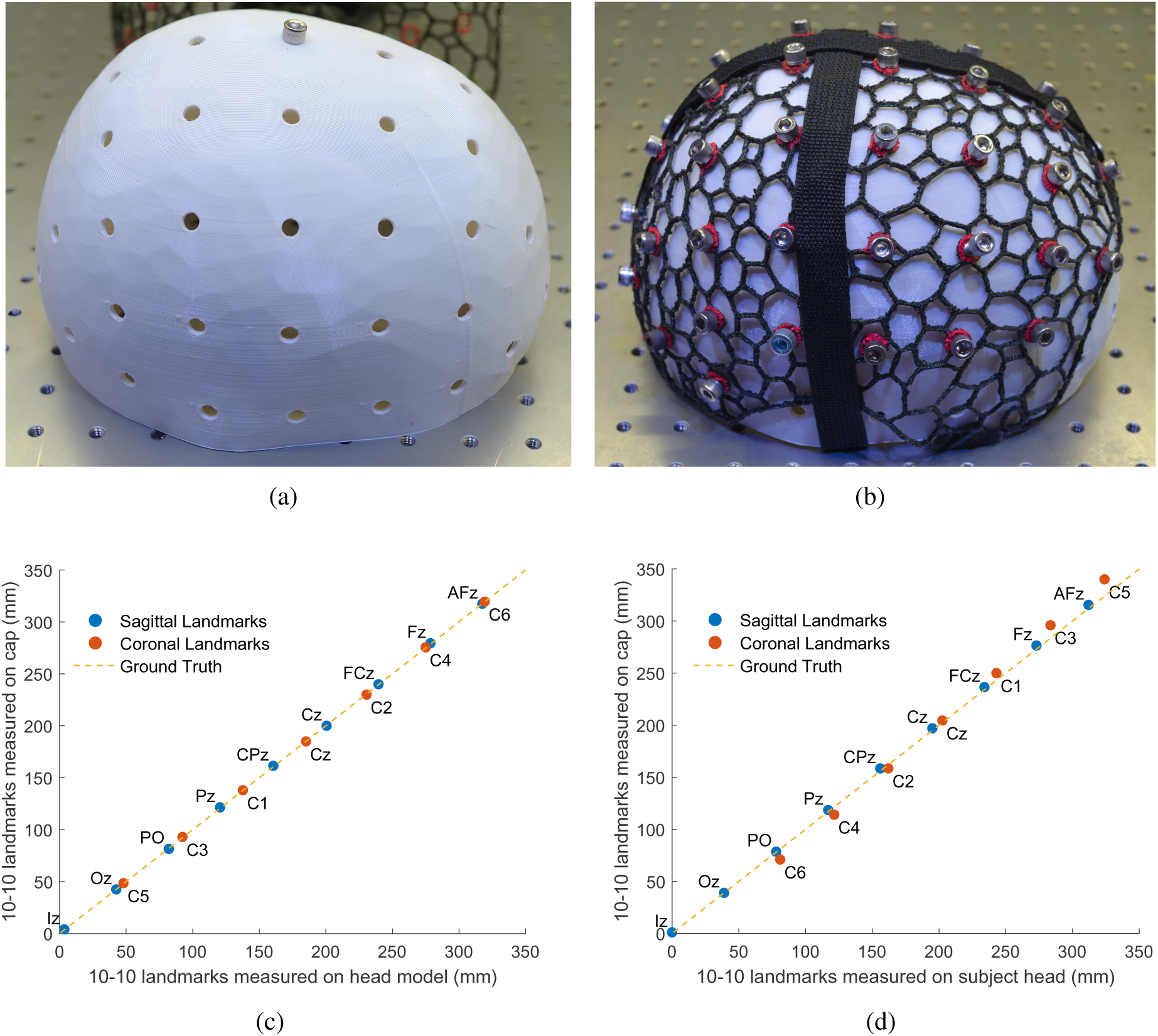
Photos of (a) a 3-D printed head model created from the NDMRI 40-44 atlas (b) donned with a head cap with pins inserted in the overlapping holes, indicating 10-20 positions. We also evaluate the accuracy of cap-embedded 10-20 positions measured over (c) a printed head model and (d) a human subject case study. Sagittal (blue circles) and coronal (red circles) landmarks are plotted. Ideally, the distances between the landmarks on the cap and head anatomy should be 1:1 (*y* = *x*), as depicted by the yellow dashed lines in (c) and (d). Each plotted point is labeled according to the 10-20 nomenclature.

Before quantifying the reproducibility of 10-10 positions on the cap, we first secure the cap over the head model by tightening two straps with one across the sagittal and one across the coronal direction. Upon simple alignment, it is clear that all 10-10 grommets on the cap are excellently aligned with the underlying through-holes over the head model. To further demonstrate the excellent and uniform spatial alignment, we place a quarter-20 sized screw across each matching pair of cap 10-10 grommet and head model through-hole. A photo showing the agreement between the cap and the head model with screws as landmark indicators is shown in Fig. 8(b).

To quantify the spatial accuracy of the printed head cap landmark locations, we measure the geodesic distances for 15*×* of the 10-10 landmark positions: Fpz, AFz, Fz, FCz, Cz, CPz, Pz, POz, and Oz on the sagittal plane (using Iz as origin), and C5, C3, C1, C2, C4 and C6 on the coronal plane (using RPA as origin)^22^ over the head model. The center of each hole is determined by averaging the positions at each edge across the diameter of the circle. Then, we place the cap over the head model and insert screws as anchoring pins at all landmarks positions that are not on the central sagittal and coronal plane. Repeating the same procedure, the distances between the origins and the centers of the circular cap grommets for each of the aforementioned 15*×* 10-10 landmarks is measured. The landmark positions on the printed head model are radially projected to the exterior surface of the cap, accounting for the radial difference due to the cap thickness. The measured distances towards the reference position (Iz or RPA) are plotted in Fig. 8(c), with the head model ground truth positions on the *x*-axis and those from the cap’s 10-10 grommet centers on the *y*-axis. We report a very strong correlation (*R*^2^ = 0.99998) and an average error of 0.4 mm *±* 0.3 mm.

#### 3.3.2 Validation on a human subject – a case study

In this case study, we aim to offer a qualitative projection on the accuracy of the 10-20 landmark grommet placements on a human subject after further considering realistic complications such as the mismatch between the subject head shape and the atlas mesh from which the cap was derived from, and the errors of the donning procedure. In this case, we use a head cap with a circumference size of 62 cm derived from the 40-44 NDMRI atlas, and place it over the head of a healthy volunteer (male, 31 years old, 61 cm head circumference, with thick dark hair). The increased circumference in the cap is desired to account for the space for hairs and the additional fNIRS assemblies.^52^ Similar to the procedure used in the above section, a total of 15*×* cap landmark positions are measured and compared against the corresponding anatomical positions derived from the volunteer’s head surface. The sagittal and coronal cranial curves are separately measured on the subject and subdivided according to the standard 10-10 measurement procedure. The inion (Iz) and right preauricular (RPA) points are designated as the origin for the sagittal and coronal curves, respectively. The cap is then positioned over the subject’s head, initially using the previously determined Cz position to guide placement. We ensure the cap is aligned along the sagittal and coronal median planes of the subject using the subject’s cranial landmarks. The subject’s head landmark distances measured along the sagittal and coronal cross sections are radially projected from the subject head surface to the outer-surface of the cap. This is achieved by applying separate geometric scaling, one for the sagittal and one for the coronal cross-section, based on the geodesic distances measured on the subject head and the outer surface of the cap. These projections allow all measured distances to be on the same surface, and also account for the elliptical shape of the head as used in other published studies.^53^ A strong correlation (*R*^2^ = 0.99893) was found between the two sets of measurements. The results are plotted in Fig. 8(d), and we report an average error of 4.7 mm *±* 4.5 mm.

### 3.4 Acquisition and validation of head cap 3-D shapes using photogrammetry

Photogrammetry has been used in DOT and fNIRS 3-D probe shape acquisitions in earlier works reported by our group^54^ as well as more recently by others.^49^ Here, we further validate the accuracy of our 3-D printed head cap 3-D shape and the recovery of the embedded 10-20 landmarks to enhance registration of optodes to the head. We apply a graphics processing unit (GPU) accelerated and machine learning-based 3-D photogrammetry software, DUST3r,^55^ and recover a full 3-D surface using photos taken around the subject’s head wearing the head cap. A total of 18 photos taken at various angles by a Pixel 2 XL Android smartphone are loaded to DUST3r’s interface. Using an NVIDIA 4090 GPU, the 3-D textured mesh model shown in Fig. 9(a) is recovered. The surface reconstruction process takes approximately one minute. This combined cap and head surface mesh model is imported to Blender and overlaid with the original 3-D cap design (gray wireframe), as shown in Fig. 9(b).

**Fig 9.**
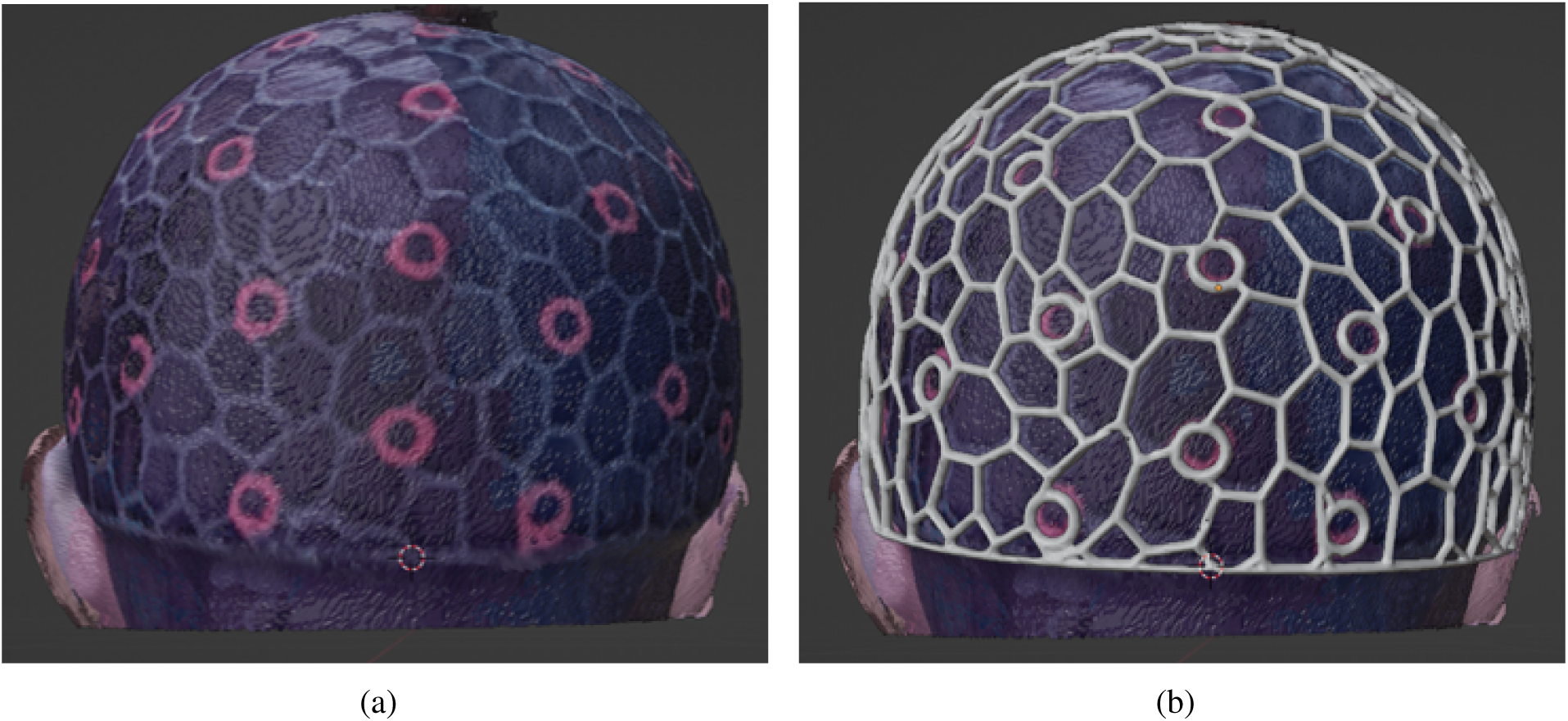
Validating cap 3-D shape accuracy using photogrammetry. In (a), we show a 3-D textured surface recovered by DUST3r photogrammetry software; the 3-D shape of the cap, including all wireframes and 10-10 markers are reconstructed along with the subject’s head surface. In (b), we import the photogrammetry recovered surface into Blender showing good alignments with the originally designed 3-D cap model (gray).

### 3.5 Utilities of 3-D printed caps in experiments

In Fig. 10, we show an example of utilizing the 3-D printed wireframe head cap in fNIRS experiments and demonstrate some of the practical considerations. In Fig. 10(a), we show a photo of a cap securing our modular optical brain imager (MOBI)^52^ over the head of a subject. In order to decide which cap to use, we first measure the circumference of the subject’s head. In addition, we also need to consider how the optical sensors are mounted to the cap. In this particular case, MOBI modules are “sandwiched” between the cap and the subject’s head,^52^ therefore, the MOBI module’s thickness, about 1 cm measured between one side of its protective silicone enclosure to the tip of the optical couplers, is multiplied by 2 and added to the head circumference. Then the cap that offers the closest match in circumference is selected. If multiple MOBI modules are used, they are first connected to each other using fixed-length ribbon cables, and then attached from inside the cap visually guided by to the cap’s built-in 10-20 markers; 3-D printed plastic locking pins and brackets (diamond shaped, shown in Fig. 10(b)) are used to quickly anchor and secure the modules over the cap wireframe. Sample *in vivo* human measurements collected using the MOBI and NeuroCaptain printed caps can be found in our recently published work.^52^

**Fig 10.**
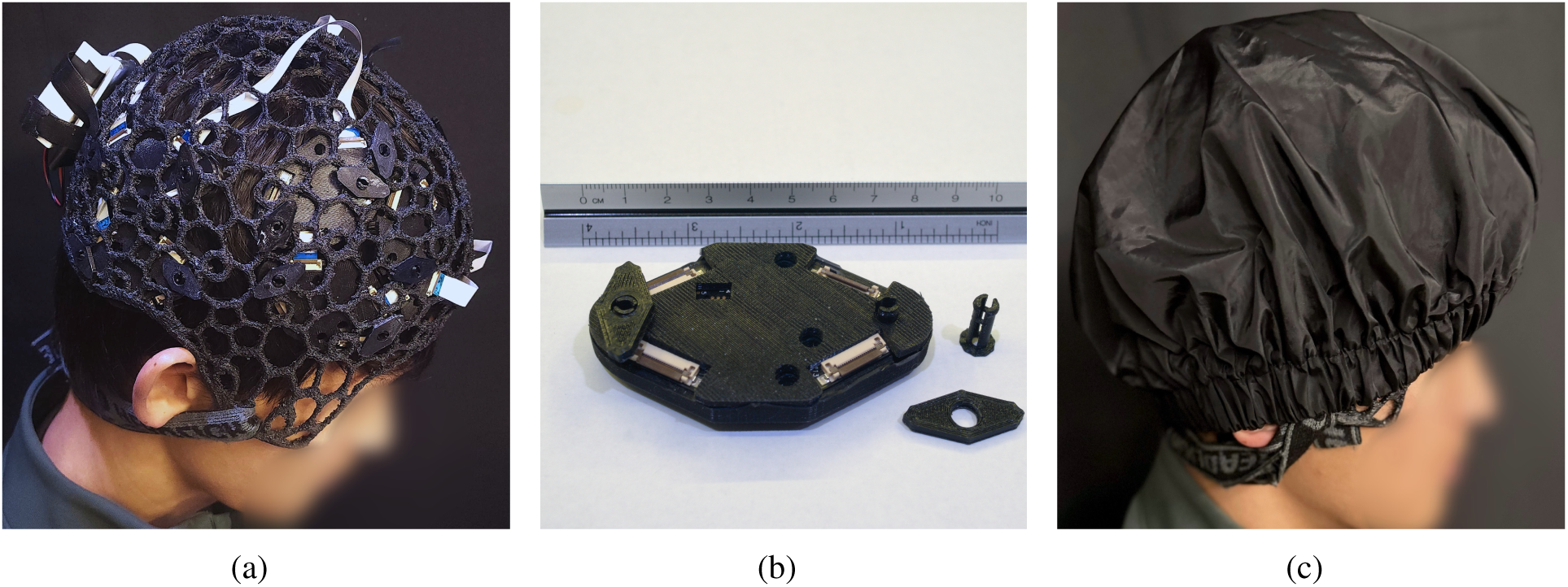
Sample application of the 3-D printed head cap in fNIRS studies, showing a) a cap carrying 10*×* modular optical brain imager (MOBI) modules placed over a subject’s head using locking pins and brackets, with a zoom-in view shown in (b), via the through-holes on the module and c) the wearable fNIRS probe enclosed by a light-blocking fabric cap. The head cap is secured by a neck strap behind the head.

Once the optical sensor units (optodes, modules etc) are secured to the cap, the full head probe is then placed over the subject’s head by first aligning the Cz grommet on the cap to the manually measured Cz position over the subject’s head and subsequently aligning the respective Nz-Iz and LPA-RPA axes. The operator can use the wireframe cap openings to access and manipulate hair around the optodes to improve optode-scalp coupling. Depending on the locations of the optodes or the experimental paradigm, a chin strap or a neck strap can be used to apply gentle tension to the optodes to ensure stable coupling. In the case shown in Fig. 10(a), a neck strap is used and conveniently pulls from the edge-piece of the cap. We found that when a task involves speaking or mouth movement, a neck strap may offer better stability compared to a chin strap. After additional adjustment in coupling and positioning, we finally enclose the entire head probe under a “shower-cap” styled light-blocking fabric cover, as shown in Fig. 10(c). In practice, the optical modules or fiber connectors can be pre-mounted to the cap to greatly shorten the setup time when the cap size does not need to change between measurements.

## 4 Discussion

Broad and easy access to standardized, low-cost and reproducible head caps^27^ is one of the key steps towards building a global community that can effectively reproduce, share and reuse neuroimaging research findings.^56^ The technologies we have developed and validated in this work, including an open-source and intuitive head cap design software pipeline, in-lab 3-D printable head caps using only off-the-shelf printers and filaments, and atlas-derived head landmark guidance are solid steps towards building such a standardized framework to achieve this goal.

A key characteristic of the caps produced from this work is the use of a wireframe design. Most commercially available fNIRS caps follow a design similar to those of EEG caps. Commonly made of a fine-mesh fabric, they serve both purposes of blocking ambient light and providing mounting position references and grommets to insert and secure optodes to the scalp. In comparison, wire-frame caps offer a number of notable ergonomic benefits. First of all, a wireframe cap provides abundant open space to access and manipulate hair underneath the cap. The presence of hair has been widely recognized as one of the top challenge in fNIRS experiments.^38^ Unfortunately, fabric based fNIRS caps enclose all hair under the cap, leaving only small openings around the optode mounting grommets to adjust hair. This often leads to inadequate adjustment. In comparison, an elastic wireframe cap has abundant open space, allowing operators to make more effective adjustments to enhance optode coupling. Secondly, the TPU material used to print the wireframe cap offers a balance between rigidity and elasticity to allow the cap to maintain a 3-D shape derived from anatomical models while having the ability to slightly deform and apply gentle pressure to the optodes. Moreover, a wireframe cap offers excellent versatility to accommodate diverse source/detector units, making it possible to rapidly mount and remount optical probes at different locations. For example, module based optodes can be secured to the wireframe via simple hooks or locking pins as shown in our MOBI module setup in Fig. 10; ferrule-shaped optodes can utilize the 3-D printed sockets or 10-20 markers to pressure-fit or lock in to the cap, or embedding additional rubber grommets at the cap’s printed grommet mounts to achieve more robust connection. In addition, a wireframe cap can also conveniently accommodate one or multiple straps to secure the cap over the subject’s head. A strap can easily loop through any of the opening near the cap edge and pull it in the desired direction. If necessary, the wireframe cap can also be easily modified by cutting additional openings or removing edges.

The use of a wireframe cap design has only recently been discussed in literature.^43^ Compared to previously reported wireframe head caps, the NeuroCaptain-derived cap offers additional benefits, namely, 1) direct fabrication in a single print, requiring no additional assembly or manual post-processing steps and thereby improving cap reproducibility, 2) retention of the 3-D anatomical shape, including all anatomically-derived and user-designated grommet/marker positions, and 3) easy customization via the intuitive Blender based interface allows users to accommodate diverse optode designs. The cap models can be conveniently 3-D printed using off-the-shelf printers and filaments. Its wireframe design also offers a semi-rigid construction that is still slightly deformable to individual-specific head surfaces while applying more uniform pressure upon optodes around the skull and better preserving the accuracy and distance between embedded 10-20 grommet locations, as demonstrated in Fig. 8(c). The primary requirement of the printer is that it must have the sufficient printing volume to accommodate the *x/y/z* dimensions of the printed cap as well as the usage of support materials to enable robust printing along the vertical *z*-direction. As a reference, the largest cap we have printed thus far has a circumference of 64 cm, with a print volume of 187 *×* 231 *×* 150 mm^3^. Regardless of these differences, both approaches carry many of the benefits of wireframe caps, as we highlighted above, and offer publicly available designers to be adopted by the broader community.

On the software side, our open-source Blender add-on, NeuroCaptain, streamlines many user-adjustable design parameters, with examples shown in Fig. 6, in an easy-to-use interface, as shown in Fig. 5. The ability to incorporate high quality head mesh generation pipelines,^45^ the support for diverse atlas models and the incorporation of quantitatively computed 10-20 system allow this toolbox to adapt to the various needs in the neuroimaging community while preserving a standardized process. The generated cap models are readily 3-D printable, and can be fabricated in-lab using off-the-shelf 3-D printers and TPU filaments. Comparing our printing experiences with the dual-filament printer (Stratasys F170) and the single-filament printer (Voron), we recommend the latter due to its lower cost, faster printing, and minimal needs for post-processing. The organic support, as shown in Fig. 7(e), offers sufficient support rigidity to maintain high 3-D accuracy and is easy to remove.

Another important point we want to raise here is that fabricating anatomically accurate head caps addresses only part of the challenges for robust and reproducible neuroimaging data acquisi-tion. To obtain reliable measurements, especially across different subjects or longitudinal measurements, the experimental operators should also pay close attention to the accuracy and consistency of donning the cap over subject’s head and placing the optodes. The spatial reposition error of fNIRS probes has only received attention recently.^57^ Incorporating built-in anatomically derived head landmarks into the head cap design provides an important visual cue for rapid and consistent probe placement over subject’s head, as well as securing optodes over the cap. Using computer vision and augmented reality to provide real-time guidance for mounting head cap and optodes, such as the work we recently reported,^58^ could supplement an anatomically-derived cap and further enhance the robustness of fNIRS studies. Regardless of donning procedures, a 3-D optode position acquisition can be applied to capture the actual mounted optode locations and compensate for the variations in positioning in post-processing. As we showed in Fig. 9(a), the rich texture in our 3-D printed cap enables fast 3-D surface acquisition using photogrammetry. Hardware based optode position estimation and tracking^52^ can also be used to improve optode placement accuracy.

We want to mention a few limitations of this current study. Firstly, the only manual step in the head cap design is the selection of cranial reference points (Nz, Iz, LPA, RPA) from the head surface mesh. The selection of LPA and RPA often depends on the accuracy of the anatomical scans or atlas models near the ear regions; the selection of Iz can also be challenging as it does not correspond to a prominent surface feature. Secondly, while NeuroCaptain streamlines the generation of 10-20 landmark grommets, one must use Blender’s built-in 3-D modeling capability to incorporate customized probe geometries and optode mounting grommets into the head cap 3-D model. This requires some user experience with Blender. Automating the process of adding user-defined probe geometries, especially transforming a 2-D flat probe representation to the head surface should be explored in future works. Thirdly, the reproducibility and accuracy of 3-D printed caps using different 3-D printers has not been characterized, and will also be investigated in the next steps.

## 5 Conclusion

In summary, we present a comprehensive neuroimaging cap design workflow, implemented as an open-source and easy-to-use Blender add-on, NeuroCaptain, to allow fNIRS and EEG researchers to conveniently create, customize, and in-house fabricate the designed caps using off-the-shelf 3-D printers and filaments. The reported cap design workflow directly capitalizes upon the powerful 3-D shape modeling capabilities from Blender, as well as a number of quantitative brain anatomical analysis toolboxes, including brain mesh generation via Brain2Mesh^45^ and Iso2Mesh,^59^ surface based 10-20 landmark estimations and JSON based data exchange formats.^44^ The printed cap features a lightweight head-conforming wireframe design, with ample open space for easy access to the hair under the cap; the built-in 10-20 head landmark grommets not only have the potential to quickly mount and secure various optical source and detector units, but also provide anatomical guidance for consistent cap donning. For easy adoption, a library of pre-generated 3-D printable cap models, derived from neurodevelopmental MRI atlas library of different age groups, are readily available for download from our website (see Data and code availability section). We showcase some of the cap designs in the Results section, and also show variations with different design settings. The software allows customization on wireframe density, thickness, 10-20 landmark density, grommet shapes, and face/ear cut-out outlines, among others. Widespread adoption of these standardized, anatomically derived head caps and freely available design tools are expected to greatly facilitate quantitative comparisons between measurements from different instruments and labs, and enable effective reuse of imaging data in secondary data analyses.

## Supporting information

Figure 5 Visualization 1

## Data and code availability

The source code of the NeuroCaptain can be downloaded at https://github.com/COTILab/NeuroCaptain; a library containing 3-D printable head cap models derived from MRI atlases of various ages can be downloaded at https://neurojson.org/db/NeuroCaptain 2024.

## Acknowledgments

This research is supported by the National Institute of Health (NIH) National Institute of Biomedical Imaging and Bioengineering grant R01-EB026998 and National Institute of Neurological Disorders and Stroke (NINDS) grant U24-NS124027.

## Footnotes

Biographies and photographs of the other authors are not available.

## References

1 G. H. Glover, “Overview of functional magnetic resonance imaging,” Neurosurgery clinics of North America 22(2), 133–139 (2011).

2 A. von Lühmann, Y. Zheng, A. Ortega-Martinez, et al., “Towards neuroscience of the every-day world (NEW) using functional near-infrared spectroscopy,” Current Opinion in Biomedical Engineering 18 (2021).

3 S. Debener, F. Minow, R. Emkes, et al., “How about taking a low-cost, small, and wireless EEG for a walk?,” Psychophysiology 49(11), 1617–1621 (2012). eprint: https://onlinelibrary.wiley.com/doi/pdf/10.1111/j.1469-8986.2012.01471.x.

4 V. Quaresima, “Functional near-infrared spectroscopy (fNIRS) for assessing cerebral cortex function during human behavior in natural/social situations: A concise review -valentina quaresima, marco ferrari, 2019,” Organizational Research Methods 22(1), 46–68 (2019).

5 E. E. Vidal-Rosas, A. v. Lühmann, P. Pinti, et al., “Wearable, high-density fNIRS and diffuse optical tomography technologies: a perspective,” Neurophotonics 10(2), 023513 (2023). Publisher: SPIE.

6 M. J. Saikia, W. G. Besio, and K. Mankodiya, “WearLight: Toward a wearable, configurable functional NIR spectroscopy system for noninvasive neuroimaging,” IEEE Transactions on Biomedical Circuits and Systems 13(1), 91–102 (2019).

7 P. Mavros, M. J Wälti, M. Nazemi, et al., “A mobile EEG study on the psychophysiological effects of walking and crowding in indoor and outdoor urban environments,” Scientific Reports 12(1), 18476 (2022). Publisher: Nature Publishing Group.

8 P. Pinti, C. Aichelburg, S. Gilbert, et al., “A review on the use of wearable functional near-infrared spectroscopy in naturalistic environments,” The Japanese Psychological Research 60(4), 347–373 (2018).

9 H. Ayaz, W. B. Baker, G. Blaney, et al., “Optical imaging and spectroscopy for the study of the human brain: status report,” Neurophotonics 9, S24001 (2022-08). Publisher: SPIE.

10 A. J. Casson, D. C. Yates, S. J. Smith, et al., “Wearable electroencephalography,” IEEE Engineering in Medicine and Biology Magazine 29(3), 44–56 (2010-05).

11 R. Li, D. Yang, F. Fang, et al., “Concurrent fNIRS and EEG for brain function investigation: A systematic, methodology-focused review,” Sensors (Basel Switzerland) 22(15), 5865 (2022-08-05).

12 K.-C. Wu, D. Tamborini, M. Renna, et al., “Open-source FlexNIRS: A low-cost, wireless and wearable cerebral health tracker,” NeuroImage 256, 119216 (2022-08-01).

13 W.-L. Chen, J. Wagner, N. Heugel, et al., “Functional near-infrared spectroscopy and its clinical application in the field of neuroscience: Advances and future directions,” Frontiers in Neuroscience 14 (2020).

14 A.-C. Ehlis, S. Schneider, T. Dresler, et al., “Application of functional near-infrared spectroscopy in psychiatry,” NeuroImage 85 **Pt** **1**, 478–488 (2014-01-15).

15 A. Bonilauri, F. Sangiuliano Intra, L. Pugnetti, et al., “A systematic review of cerebral functional near-infrared spectroscopy in chronic neurological diseases—actual applications and future perspectives,” Diagnostics 10(8), 581 (2020-08-12).

16 L. K. Butler, S. Kiran, and H. Tager-Flusberg, “Functional near-infrared spectroscopy in the study of speech and language impairment across the life span: A systematic review,” American Journal of Speech-Language Pathology 29(3), 1674–1701 (2020-08).

17 Y. Li, L. Zhang, K. Long, et al., “Real-time monitoring prefrontal activities during misc video game playing by functional near-infrared spectroscopy,” Journal of Biophotonics 11(9), e201700308 (2018). eprint: https://misclibrary.wiley.com/doi/pdf/10.1002/jbio.201700308.

18 A. Girouard, E. T. Solovey, and R. J. Jacob, “Designing a passive brain computer interface using real time classification of functional near-infrared spectroscopy,” International Journal of Autonomous and Adaptive Communications Systems 6(1), 26 (2013).

19 M. A. Yücel, J. Selb, D. A. Boas, et al., “Reducing motion artifacts for long-term clinical NIRS monitoring using collodion-fixed prism-based optical fibers,” NeuroImage 85, 192– 201 (2014-01-15).

20 A. Kassab, J. L. Lan, P. Vannasing, et al., “Functional near-infrared spectroscopy caps for brain activity monitoring: a review,” Applied Optics 54(3), 576–586 (2015).

21. 21 S. A. Khezri, “The influence of optode pressure on the quality of functional near-infrared spectroscopy signal,” (2020).

22 R. Oostenveld and P. Praamstra, “The five percent electrode system for high-resolution EEG and ERP measurements,” Clinical Neurophysiology 112 (2001).

23 V. Jurcak, D. Tsuzuki, and I. Dan, “10/20, 10/10, and 10/5 systems revisited: their validity as relative head-surface-based positioning systems,” NeuroImage 34(4), 1600–1611 (2007-02-15).

24 P. Giacometti, K. L. Perdue, and S. G. Diamond, “Algorithm to find high density EEG scalp coordinates and analysis of their correspondence to structural and functional regions of the brain,” Journal of Neuroscience Methods 229, 84–96 (2014-05-30).

25 H. Sherkat, T. Gjvaag, and P. Mirtaheri, “Experimental investigation on the light transmission of a textile-based over-cap used in functional near-infrared spectroscopy,” Clinical and Preclinal Optical Diagnostics II **E****B101** (2019).

26 A. Kassab and M. Sawan, “The NIRS cap: Key part of emerging wearable brain-device interfaces,” in Developments in Near-Infrared Spectroscopy, K. G. Kyprianidis and J. Skvaril, Eds., InTech (2017-03-15).

27 A. v. Lühmann, “Can the fNIRS community design a standard cap layout for uniform whole-head HD fNIRS coverage? a discussion.,” in The Society for functional Near Infrared Spectroscopy, (2022-01-01).

28 F. Tsow, A. Kumar, S. H. Hosseini, et al., “A low-cost, wearable, do-it-yourself functional near-infrared spectroscopy (DIY-fNIRS) headband,” HardwareX 10, e00204 (2021-10-01).

29 A. von Lühmann, C. Herff, D. Heger, et al., “Toward a wireless open source instrument: Functional near-infrared spectroscopy in mobile neuroergonomics and BCI applications,” Frontiers in Human Neuroscience 9, 153157 (2015-11-12). Publisher: Frontiers.

30 P.-Y. Lin, N. Roche-Labarbe, M. Dehaes, et al., “Non-invasive optical measurement of cerebral metabolism and hemodynamics in infants,” Journal of Visualized Experiments : JoVE 73(73), 4379 (2013-03-14).

31 B. R. White and J. P. Culver, “Quantitative evaluation of high-density diffuse optical tomography: in vivo resolution and mapping performance,” Journal of Biomedical Optics 15(2), 026006 (2010-03). Publisher: SPIE.

32 C. L. Scrivener and A. T. Reader, “Variability of EEG electrode positions and their underlying brain regions: visualizing gel artifacts from a simultaneous EEG-fMRI dataset,” Brain and Behavior 12(2), e2476 (2022).

33 S. R. Atcherson, H. J. Gould, M. A. Pousson, et al., “Variability of electrode positions using electrode caps,” Brain Topography 20(2), 105–111 (2007-12-01).

34 S. L. Novi, E. J. Forero, J. A. I. Rubianes Silva, et al., “Integration of spatial information increases reproducibility in functional near-infrared spectroscopy,” Frontiers in Neuroscience 14 (2020).

35 S. Srinivasan, D. Acharya, E. Butters, et al., “Subject-specific information enhances spatial accuracy of high-density diffuse optical tomography,” Frontiers in Neuroergonomics 5 (2024-02-19). Publisher: Frontiers.

36 S. Brigadoi, D. Salvagnin, M. Fischetti, et al., “Array designer: automated optimized array design for functional near-infrared spectroscopy,” Neurophotonics 5(3), 035010 (2018-07).

37 F. Klein, “Optimizing spatial specificity and signal quality in fNIRS: an overview of potential challenges and possible options for improving the reliability of real-time applications,” Frontiers in Neuroergonomics 5 (2024). Publisher: Frontiers.

38 B. Khan, C. Wildey, R. Francis, et al., “Improving optical contact for functional near-infrared brain spectroscopy and imaging with brush optodes,” Biomedical Optics Express 3(5), 878– 898 (2012-04-06).

39 E. M. Frijia, A. Billing, S. Lloyd-Fox, et al., “Functional imaging of the developing brain with wearable high-density diffuse optical tomography: A new benchmark for infant neuroimaging outside the scanner environment,” Neuroimage 225, 117490 (2021-01-15).

40 J. M. Baker, J. L. Bruno, A. Gundran, et al., “fNIRS measurement of cortical activation and functional connectivity during a visuospatial working memory task,” PLoS ONE 13(8), e0201486 (2018-08-02).

41 A. Kassab, J. Le Lan, J. Tremblay, et al., “Multichannel wearable fNIRS-EEG system for long-term clinical monitoring,” Human Brain Mapping 39(1), 7–23 (2018-01).

42 H. Matsumura, T. Tanijiri, M. Kouchi, et al., “Global patterns of the cranial form of modern human populations described by analysis of a 3d surface homologous model,” Scientific Reports 12(1), 13826 (2022-08-15). Number: 1 Publisher: Nature Publishing Group.

43 A. v. Lühmann, S. Kura, W. J. O’Brien, et al., “ninjaCap: a fully customizable and 3D printable headgear for functional near-infrared spectroscopy and electroencephalography brain imaging,” Neurophotonics 11(3), 036601 (2024). Publisher: SPIE.

44 Y. Zhang and Q. Fang, “BlenderPhotonics: an integrated open-source software environ-ment for three-dimensional meshing and photon simulations in complex tissues,” Journal of Biomedical Optics 27(8), 083014 (2022-08).

45 A. P. Tran, S. Yan, and Q. Fang, “Improving model-based functional near-infrared spectroscopy analysis using mesh-based anatomical and light-transport models,” Neurophotonics 7(1), 015008 (2020-01).

46 M. Jenkinson, C. F. Beckmann, T. E. J. Behrens, et al., “FSL,” NeuroImage 62(2), 782–790 (2012-08-15).

47 D. W. Shattuck and R. M. Leahy, “BrainSuite: an automated cortical surface identification tool,” Medical Image Analysis 6(2), 129–142 (2002-06).

48 R. J. Cooper, M. Caffini, J. Dubb, et al., “Validating atlas-guided DOT: a comparison of diffuse optical tomography informed by atlas and subject-specific anatomies,” NeuroImage 62(3), 1999–2006 (2012-09).

49 I. Mazzonetto, M. Castellaro, R. J. Cooper, et al., “Smartphone-based photogrammetry provides improved localization and registration of scalp-mounted neuroimaging sensors,” Scientific Reports 12(1), 10862 (2022). Publisher: Nature Publishing Group.

50 J.-D. Boissonnat and S. Oudot, “Provably good sampling and meshing of surfaces,” Graphical Models 67, 405–451 (2005).

51 M. S. Myslobodsky, R. Coppola, J. Bar-Ziv, et al., “Adequacy of the international 10-20 electrode system for computed neurophysiologic topography,” Journal of Clinical Neurophysiology: Official Publication of the American Electroencephalographic Society 7(4), 507–518 (1990).

52 E. Xu, M. Vanegas, M. Mireles, et al., “Flexible circuit-based spatially aware modular optical brain imaging system for high-density measurements in natural settings,” Neurophotonics 11(3), 035002 (2024).

53 X. Zhai, H. Santosa, and T. J. Huppert, “Using anatomically defined regions-of-interest to adjust for head-size and probe alignment in functional near-infrared spectroscopy,” Neurophotonics 7(2) (2020-09-23).

54 Q. Fang, “Quantitative diffuse optical tomography using a mobile phone camera and automatic 3D photo stitching,” Biomedical Optics and 3-D Imaging BSu3A, BSu3A.96, Optica Publishing Group (2012).

55 S. Wang, V. Leroy, Y. Cabon, et al., “DUSt3R: Geometric 3d vision made easy,” (2023).

56 M. A. Yücel, A. v. Lühmann, F. Scholkmann, et al., “Best practices for fNIRS publications,” Neurophotonics 8(1), 012101 (2021-01). Publisher: SPIE.

57 A. Blasi, S. Lloyd-Fox, M. H. Johnson, et al., “Test–retest reliability of functional near infrared spectroscopy in infants,” Neurophotonics 1(2), 025005 (2014-09). Publisher: SPIE.

58 F.-Y. Yen, Y.-A. Lin, and Q. Fang, “Real-time guidance for fNIRS headgear placement using augmented reality,” Optica Biophotonics Congress: Biomedical Optics 2024 Optics and the Brain (BRAIN), BW1C.6, Optica Publishing Group (2024).

59 Q. Fang and D. A. Boas, “Tetrahedral mesh generation from volumetric binary and gray-scale images,” in Proceedings of the Sixth IEEE international conference on Symposium on Biomedical Imaging: From Nano to Macro, ISBI’09, 1142–1145, IEEE Press (2009).

